# KrakenHLL: Confident and fast metagenomics classification using unique k-mer counts

**DOI:** 10.1101/262956

**Authors:** FP Breitwieser, SL Salzberg

## Abstract

False positive identifications are a significant problem in metagenomic classification. We present KrakenHLL, a novel metagenomic classifier that combines the fast k-mer based classification of Kraken with an efficient algorithm for assessing the coverage of unique k-mers found in each species in a dataset. On various test datasets, KrakenHLL gives better recall and F1-scores than other methods, and effectively classifies and distinguishes pathogens with low abundance from false positives in infectious disease samples. By using the probabilistic cardinality estimator HyperLogLog (HLL), KrakenHLL is as fast as Kraken and requires little additional memory.

## Background

Metagenomic classifiers attempt to assign a taxonomic identity to each read in a data set. Because metagenomics data often contain tens of millions of reads, classification is typically done using exact matching of short words of length *k* (*k*-mers) rather than alignment, which would be unacceptably slow. The results contain read classifications but not their aligned positions in the genomes [as reviewed by 1]. However, read counts can be deceiving. Sequence contamination of the samples–introduced from laboratory kits or the environment during sample extraction, handling or sequencing–can yield high numbers of spurious identifications [2, 3]. Having only small amounts of input material can further compound the problem of contamination. When using sequencing for clinical diagnosis of infectious diseases, for example, less than 0.1% of the DNA may derive from microbes of interest [4, 5]. Additional spurious matches can result from low-complexity regions of genomes and from contamination in the database genomes themselves [6].

Such false-positive reads typically match only small portions of a genome; e.g., if a species’ genome contains a low-complexity region, and the only reads matching that species fall in this region, then the species was probably not present in the sample. Reads from microbes that are truly present should distribute relatively uniformly across the genome rather than being concentrated in one or a few locations. Genome alignment can reveal this information. However, alignment is resource intensive, requiring the construction of indexes for every genome and a relatively slow alignment step to compare all reads against those indexes. Some metagenomics methods do use coverage information to improve mapping or quantification accuracy, but these methods require results from much slower alignment methods as input [7]. Assembly-based methods also help to avoid false positives, but these are useful only for highly abundant species [8].

Here, we present KrakenHLL, a novel method that combines very fast k-mer based classification with a fast k-mer cardinality estimation. KrakenHLL is based on the Kraken metagenomics classifier [9], to which it adds a method for counting the number of unique k-mers identified for each taxon using the efficient cardinality estimation algorithm HyperLogLog [10-12]. By counting how many of each genome’s unique k-mers are covered by reads, KrakenHLL can often discern false positive from true positive matches. Furthermore, KrakenHLL implements additional new features to improve metagenomics classification: (a) searches can be done against multiple databases hierarchically, (b) the taxonomy can be extended to include nodes for strains and plasmids, thus enabling their detection, and (c) the database build script allows the addition of >100,000 viruses from the NCBI Viral Genome Resource [13]. KrakenHLL provides a superset of the information provided by Kraken while running equally fast or slightly faster, and while using very little additional memory during classification.

## Results

KrakenHLL was developed to provide efficient k-mer count information for all taxa identified in a metagenomics experiment. The main workflow is as follows: As reads are processed, each k-mer is assigned a taxon from the database (Figure 1A). KrakenHLL instantiates a HyperLogLog data sketch for each taxon, and adds the k-mers to it (Figure 1B and Supplementary Information). After classification of a read, KrakenHLL traverses up the taxonomic tree and merges the estimators of each taxon with its parent. In its classification report, KrakenHLL includes the number of unique k-mers and the depth of k-mer coverage for each taxon that it observed in the input data (Figure 1C).

**Figure 1.**
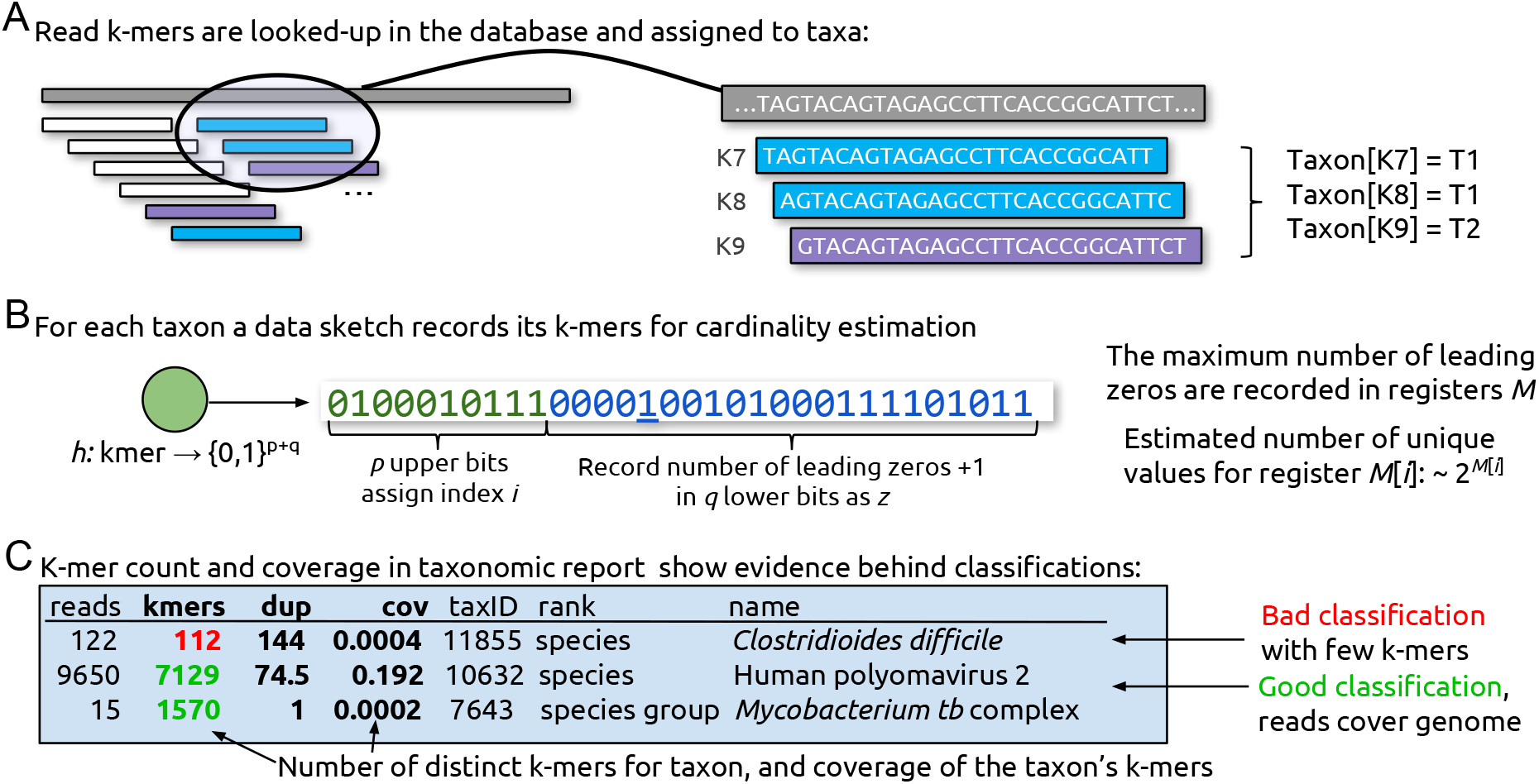
Overview of the KrakenHLL algorithm and output. (A) An input read is shown as a long gray rectangle, with k-mers shown as shorter rectangles below it. The taxon mappings for each k-mer are compared to the database, shown as larger rectangles on the right. For each taxon, a unique k-mer counter is instantiated, and the observed k-mers (K7, K8, and K9) are added to the counters. (B) Unique k-mer counting is implemented with the probabilistic estimation method HyperLogLog (HLL) using 16KB of memory per counter, which limits the error in the cardinality estimate to 1% (see main text). (C) The output includes the number of reads, unique k-mers, duplicity (average time each k-mer has been seen) and coverage for each taxon observed in the input data.

### Efficient k-mer cardinality estimation using the HyperLogLog algorithm

Cardinality is the number of elements in a set without duplicates; *e. g.* the number of distinct words in a text. An exact count can be kept by storing the elements in a sorted list or linear probing hash table, but that requires memory proportional to the number of unique elements. When an accurate estimate of the cardinality is sufficient, however, the computation can be done efficiently with very small amount of fixed memory. The HyperLogLog algorithm (HLL) [10], which is well suited for k-mer counting [14], keeps a summary or *sketch* of the data that is sufficient for precise estimation of the cardinality, but requires only a small amount of constant space to estimate cardinalities up to billions. The method centers on the idea that long runs of leading zeros, which can be efficiently computed using machine instructions, are unlikely in random bitstrings. For example, about every fourth bitstring in a random series should start with 01_2_ (one 0-bit before the first 1-bit), and about every 32^nd^ hash starts with 00001_2_. Conversely, if we know the maximum number of leading zeros *k* of the members of a random set, we can use 2*^k+1^* as a crude estimate of its cardinality (more details in the Suppl. Methods). HLL keeps *m*=2*^p^* one-byte counts of the maximum numbers of leading zeros on the data (its data *sketch*), with *p*, the precision parameter, typically between 10 and 18 (see Figure 2). For cardinalities up to *m*/4, we use the sparse representation of the registers suggested by Heule et al. [11] that has the much higher effective precision *p’* of 25 by encoding each index and count in a vector of four-byte values. To add a k-mer to its taxon’s sketch, the k-mer (with k up to 31) is first mapped by a hash function to a 64-bit hash value. Note that k-mers that contain non-A, C, G or T characters (such as ambiguous IUPAC characters) are ignored by KrakenHLL. The first *p* bits of the hash value are used as index *i*, and the later 64-*p*=*q* bits for counting the number of leading zeros *k*. The value of the register *M[i]* in the sketch is updated if *k* is larger than the current value of *M[i]*.

When the read classification is finished, the taxon sketches are aggregated up the taxonomy tree by taking the maximum of each register value. The resulting sketches are the same as if the k-mers were counted at their whole lineage from the beginning. KrakenHLL then computes cardinality estimates using the formula proposed by Ertl [12], which has theoretical and practical advantages and does not require empirical bias correction factors [10, 11]. In our tests it performed better than Flajolet’s and Heule’s methods (Suppl. Figures 1 and 2).

The expected relative error of the final cardinality estimate is approximately 1.04/sqrt(*2^p^*) [10]. With *p*=14, the sketch uses 2^14^ one-byte registers, i.e. 16 KB of space, and gives estimates with relative errors of less than 1% (Figure 2). An exact counter would require about 40 MB per million distinct k-mers when implemented using an unordered set; i. e. about 40 GB for the pathogen identification samples with an average of one billion distinct k-mers per sample. However, unordered sets have worst case insertion time complexity linear to the container size (and require re-hashes on resize), while it is constant for HLL.

| $p$ | $m=2^p$ | Space<br>(kB) | Rel.<br>Error |
| --- | --- | --- | --- |
| 10 | 1024 | 1 | 3.25% |
| 11 | 2048 | 2 | 2.23% |
| 12 | 4096 | 4 | 1.63% |
| 13 | 8192 | 8 | 1.15% |
| 14 | 16384 | 16 | 0.81% |
| 15 | 32768 | 32 | 0.57% |
| 16 | 65536 | 64 | 0.41% |
| 17 | 131072 | 128 | 0.29% |
| 18 | 262144 | 256 | 0.20% |
| 25 |  |  | 0.02% |

**Figure 2:**
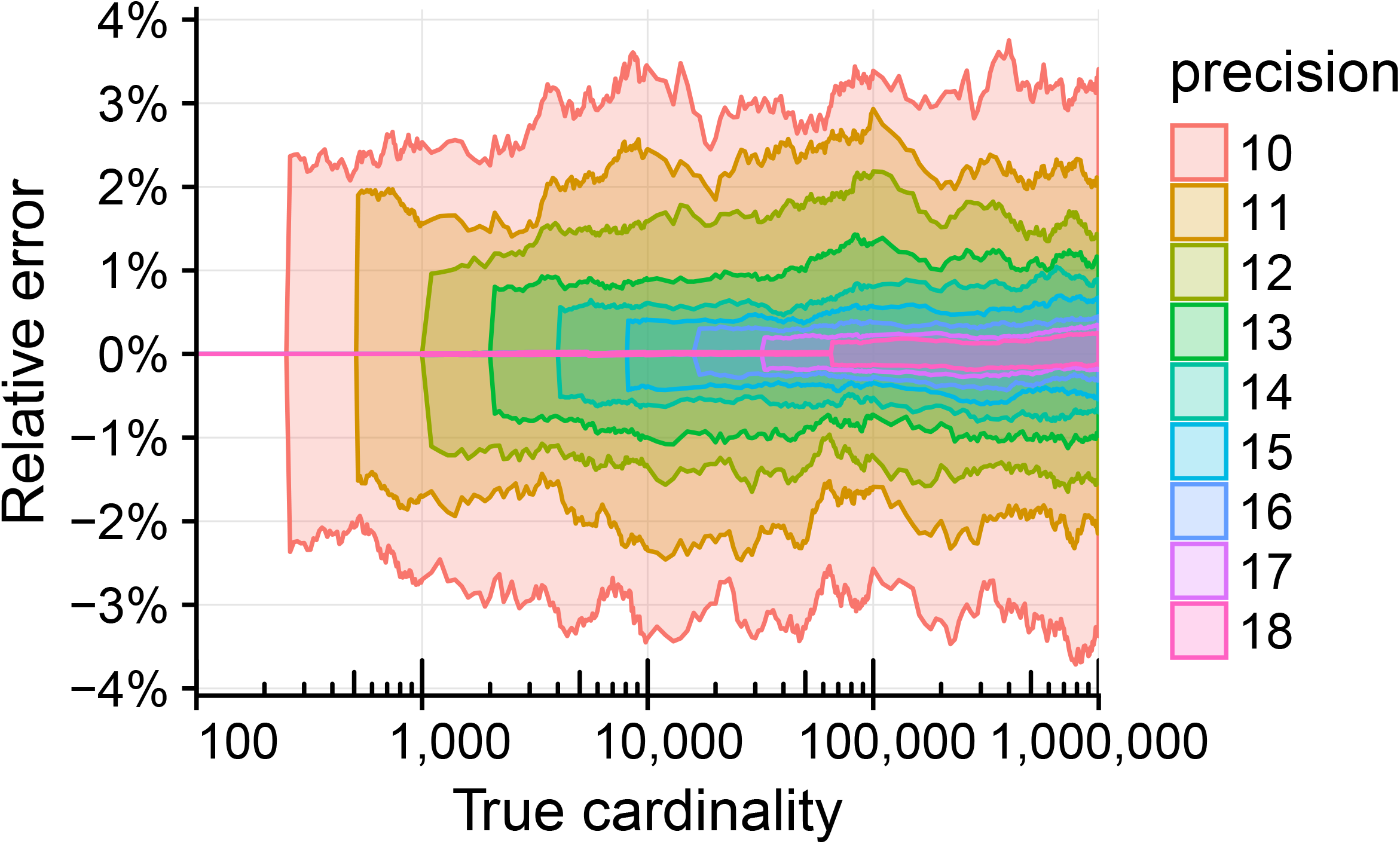
Cardinality estimation using HyperLogLog for randomly sampled k-mers from microbial genomes. Left: standard deviations of the relative errors of the estimate with precision *p* ranging from 10 to 18. No systematic biases are apparent, and, as expected, the errors decrease with higher values of *p*. Up to cardinalities of about 2^*p*^/4, the relative error is near zero. At higher cardinalities, the error boundaries stay near constant. Right: the size of the registers, space requirement, and expected relative error for HyperLogLog cardinality estimates with different values of *p*. For example, with a precision *p*=14, the expected relative error is 0.81% and the counter only requires 16 KB of space, which is three orders of magnitude less than that of an exact counter (at a cardinality of one million). Up to cardinalities of 2^*p*^/4, KrakenHLL uses a sparse representation of the counter with a higher precision of 25 and an effective relative error rate of about 0.02%.

### Results on twenty-one simulated and ten biological test datasets

We assessed KrakenHLL’s performance on the 34 datasets compiled by McIntyre et al. [15] (see Suppl. Table 3). We place greater emphasis on the eleven biological datasets, which contain more realistic laboratory and environmental contamination. In the first part of this section, we show that unique k-mer counts provide higher classification accuracy than read counts, and in the second part we compare KrakenHLL with the results of eleven metagenomics classifiers. We ran KrakenHLL on three databases: ‘orig’, the database used by McIntyre et al., ‘std’, which contains all current complete bacterial, archaeal and viral genomes from RefSeq plus viral neighbor sequences and the human reference genome, and ‘nt’, which contains all microbial sequences (including fungi and protists) in the non-redundant nucleotide collection nr/nt provided by NCBI (see Suppl. Methods Section 2 for details). The ‘std’ database furthermore includes the UniVec and EmVec sequence sets of synthetic constructs and vector sequences; and low-complexity k-mers in microbial sequences were masked using NCBI’s dustmasker with default settings. We use two metrics to compare how well methods can separate true positives and false positives: (a) F1 score, i. e. the harmonic mean of precision *p* and recall *r*, and (b) recall at a maximum false discovery rate (FDR) of 5%. For each method, we compute and select the ideal thresholds based on the read count, k-mer count or abundance calls. Precision *p* is defined as the number of correctly called species (or genera) divided by the number of all called species (or genera) at a given threshold. Recall *r* is the proportion of species (or genera) that are in the test dataset and that are called at a given threshold. Higher F1 scores indicate a better separation between true positives and false positives. Higher recall means that more true species can be recovered while controlling the false positives.

Because the NCBI taxonomy has been updated since the datasets were published, we manually updated the “truth” sets in several datasets (see Suppl. Methods Section 2.3 for all changes). Any cases that might have been missed would result in a lower apparent performance of KrakenHLL. Note that we exclude the over ten-year-old simulated datasets simHC, simMC and simLC from Mavromatis et al. (2007), as well as the biological dataset JGI SRR033547 which has only 100 reads.

**Table 1:**
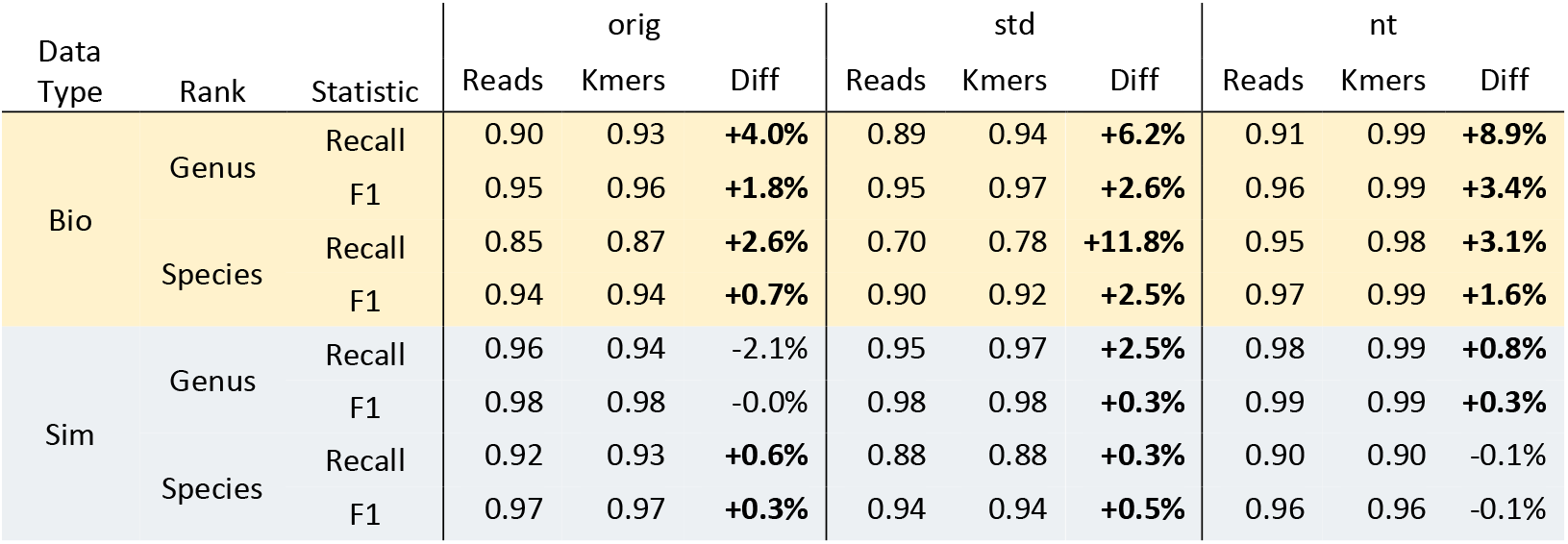
Performance of read count and unique k-mer thresholds on 10 biological and 21 simulated datasets against three databases (‘orig’, ‘std’, ‘nt). Unique k-mer count thresholds give up to 10% better recall and F1 scores, particularly for the biological datasets.

| Data Type | Rank | Statistic | orig |  |  | std |  |  | nt |  |  |
| --- | --- | --- | --- | --- | --- | --- | --- | --- | --- | --- | --- |
|  |  |  | Reads | Kmers | Diff | Reads | Kmers | Diff | Reads | Kmers | Diff |
| Bio | Genus | Recall | 0.90 | 0.93 | **+4.0%** | 0.89 | 0.94 | **+6.2%** | 0.91 | 0.99 | **+8.9%** |
|  |  | F1 | 0.95 | 0.96 | **+1.8%** | 0.95 | 0.97 | **+2.6%** | 0.96 | 0.99 | **+3.4%** |
|  | Species | Recall | 0.85 | 0.87 | **+2.6%** | 0.70 | 0.78 | **+11.8%** | 0.95 | 0.98 | **+3.1%** |
|  |  | F1 | 0.94 | 0.94 | **+0.7%** | 0.90 | 0.92 | **+2.5%** | 0.97 | 0.99 | **+1.6%** |
| Sim | Genus | Recall | 0.96 | 0.94 | -2.1% | 0.95 | 0.97 | **+2.5%** | 0.98 | 0.99 | **+0.8%** |
|  |  | F1 | 0.98 | 0.98 | -0.0% | 0.98 | 0.98 | **+0.3%** | 0.99 | 0.99 | **+0.3%** |
|  | Species | Recall | 0.92 | 0.93 | **+0.6%** | 0.88 | 0.88 | **+0.3%** | 0.90 | 0.90 | -0.1% |
|  |  | F1 | 0.97 | 0.97 | **+0.3%** | 0.94 | 0.94 | **+0.5%** | 0.96 | 0.96 | -0.1% |

#### Unique k-mer versus read count thresholds

We first looked at the performance of the unique k-mer count thresholds versus read count thresholds (as would be used with Kraken). The k-mer count thresholds worked very well, particularly for the biological datasets (Table 1 and Suppl. Table 3). On the genus level, the average recall in the biological datasets increases by 4-9%, and the average F1 score increases 2-3%. On the species level, the average increase in recall in the biological sets is between 3 and 12%, and the F1 score increases by 1-2%.

On the simulated datasets, the differences are less pronounced and vary between databases, even though on average the unique k-mer count is again better. However, only in two cases (genus recall on databases ‘orig’ and ‘std’) the difference is higher than 1% in any direction. We find that simulated datasets often lack false positives with a decent number of reads but a lower number of unique k-mer counts, which we see in real data. Instead, in most simulated datasets the number of unique k-mers is linearly increasing with the number of unique reads in both true and false positives (Suppl. Figure 4). In biological datasets, sequence contamination and lower read counts for the true positives make the task of separating true and false positives harder.

#### Comparison of KrakenHLL with eleven other methods

Next, we compared KrakenHLL’s unique k-mer counts with the results of eleven metagenomics classifiers from McIntyre et al. [15], which include the alignment-based methods Blast + Megan [16, 17], Diamond + Megan [17, 18] and MetaFlow [19], the k-mer based CLARK [20], CLARK-S [21], Kraken [9], LMAT [22], NBC [23] and the marker-based methods GOTTCHA [24], MetaPhlAn2 [25], PhyloSift [26]. KrakenHLL with database ‘nt’ has the highest average recall and F1 score across the biological datasets, as shown in Table 2. As seen before, using unique k-mer instead of read counts as thresholds increases the scores. While the database selection proves to be very important (KrakenHLL with database ‘std’ is performing 10% worse than KrakenHLL with database ‘nt’), only Blast has higher average scores than KrakenHLL with k-mer count thresholds on the original database. On the simulated datasets, KrakenHLL with the ‘nt’ database still ranks at the top, though, as seen previously there is more variation (Suppl. Table 4). Notably CLARK is as good as KrakenHLL, but Blast has much worse scores on the simulated datasets.

**Table 2:** Performance of KrakenHLL (with unique k-mer count thresholds) compared to metagenomic classifiers [15] on the biological datasets (n=10). F1 and Recall show the average values over the datasets. Note that ‘KrakenHLL reads’ would be equivalent to standard Kraken.

|  | Genus |  | Species |  | Avg |
| --- | --- | --- | --- | --- | --- |
|  | F1 | Recall | F1 | Recall |  |
| KrakenHLL nt kmers | **0.99** | **0.99** | **0.99** | **0.98** | **0.99** |
| KrakenHLL nt reads | 0.96 | 0.91 | 0.97 | 0.95 | 0.95 |
| BlastMeganFilteredLiberal | 0.97 | 0.94 | 0.97 | 0.89 | 0.94 |
| BlastMeganFiltered | 0.97 | 0.93 | 0.96 | 0.87 | 0.93 |
| KrakenHLL orig kmers | 0.96 | 0.93 | 0.94 | 0.87 | 0.93 |
| ClarkM4Spaced | 0.95 | 0.90 | 0.94 | 0.88 | 0.92 |
| KrakenHLL orig reads | 0.95 | 0.90 | 0.94 | 0.85 | 0.91 |
| Kraken | 0.95 | 0.90 | 0.94 | 0.84 | 0.91 |
| KrakenHLL std kmers | 0.97 | 0.94 | 0.92 | 0.78 | 0.90 |
| DiamondMegan_sensitive | 0.98 | 0.93 | 0.92 | 0.74 | 0.89 |
| KrakenFiltered | 0.95 | 0.91 | 0.90 | 0.75 | 0.88 |
| ClarkM1Default | 0.94 | 0.85 | 0.91 | 0.77 | 0.87 |
| KrakenHLL std reads | 0.95 | 0.89 | 0.90 | 0.70 | 0.86 |
| LMAT | 0.97 | 0.93 | 0.91 | 0.60 | 0.85 |
| DiamondMegan | 0.94 | 0.87 | 0.91 | 0.66 | 0.85 |
| Gottcha | 0.91 | 0.84 | 0.87 | 0.67 | 0.82 |
| NBC | 0.87 | 0.76 | 0.85 | 0.73 | 0.80 |
| Metaphlan | 0.94 | 0.89 | 0.83 | 0.55 | 0.80 |
| MetaFlow | 0.66 | 0.53 | 0.65 | 0.51 | 0.59 |
| PhyloSift | 0.68 | 0.29 | 0.78 | 0.54 | 0.57 |
| PhyloSift90pct | 0.68 | 0.30 | 0.77 | 0.52 | 0.57 |

### Generating a better test dataset, and selecting an appropriate k-mer threshold

In the previous section we demonstrated that KrakenHLL gives better recall and F1-scores than other classifiers on the test datasets, given the correct thresholds. How can the correct thresholds be determined on real data with varying sequencing depths and complex communities? The test datasets are not ideal for that: The biological datasets lack complexity with a maximum of 25 species in some of the samples, while the simulated samples lack the features of biological datasets.

We thus generated a third type of test dataset by sampling reads from real bacterial isolate sequencing runs, of which there are tens of thousands in the Sequence Read Archive (SRA). That way we created a complex test dataset for which we know the ground truth, with all the features of real sequencing experiments, including lab contaminants and sequencing errors. We selected 280 SRA datasets from 280 different bacterial species that are linked to complete RefSeq genomes (see Suppl. Methods Section 2.4). We randomly sampled between one hundred and one million reads (logarithmically distributed) from each experiment, which gave 34 million read pairs in total. Furthermore, we sub-sampled five read sets with between one to twenty million reads. All read sets were classified with KrakenHLL using the ‘std’ database.

**Figure 3:**
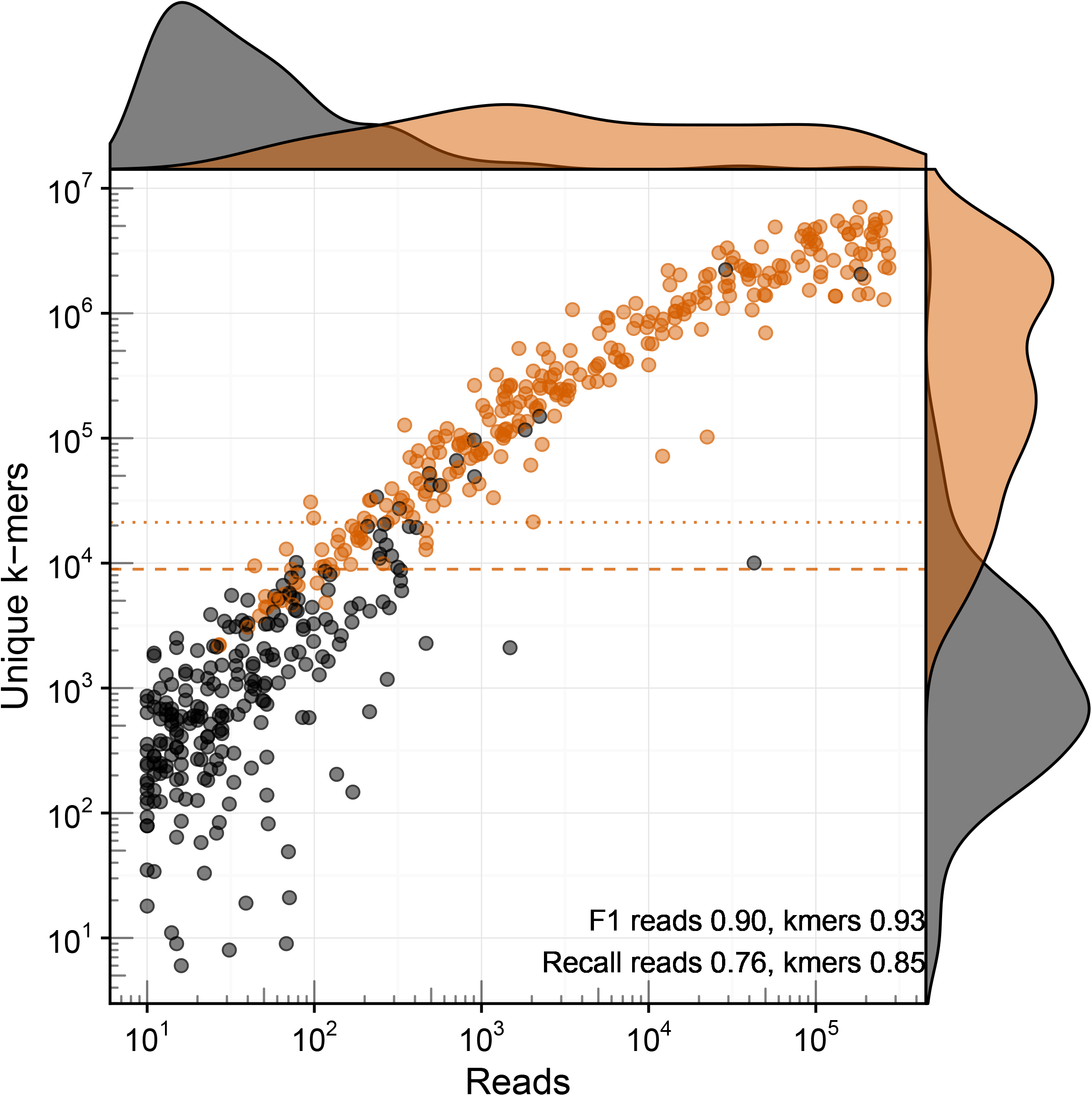
Unique k-mer count separates true and false positives better than read counts in a complex dataset with ten million reads sampled from SRA experiments. Each dot represents a species, with true species in orange and false species in black. The dashed and dotted lines show the k-mer thresholds for the ideal F1 score and recall at a maximum of 5% FDR, respectively. In this dataset, a unique k-mer count in the range 10000–20000 would give the best threshold for selecting true species.

Consistent with the results of the previous section, we found that unique k-mer counts provide better thresholds than read counts both in terms of F1 score and recall in all test datasets (e.g. Figure 3 on ten million reads – species recall using k-mers is 0.85, recall using reads 0.76). With higher sequencing depth, the recall increased slightly - from 0.80 to 0.85 on the species level, and from 0.87 to 0.89 on the genus level. The ideal values of the unique k-mer count thresholds, however, vary widely with different sequencing depths. We found that the ideal thresholds increase by about 2000 unique k-mers per one million reads (see Figure 4). McIntyre et al. [15] found that k-mer based methods show a positive relationship between sequencing depths and misclassified reads. Our analysis also shows that with deeper sequencing depths higher thresholds are required to control the false-positive rate.

| No. of reads | Fraction | Genus |  | Species |  |
| --- | --- | --- | --- | --- | --- |
|  |  | Threshold | Recall | Threshold | Recall |
| 1 million | 0.03 | 2555 | 0.87 | 3682 | 0.80 |
| 2 million | 0.06 | 4483 | 0.86 | 6152 | 0.81 |
| 5 million | 0.15 | 12723 | 0.87 | 10459 | 0.85 |
| 10 million | 0.3 | 21896 | 0.88 | 21201 | 0.85 |
| 20 million | 0.6 | 43417 | 0.88 | 43417 | 0.84 |
| 34.3 million | 1 | 69847 | 0.89 | 688842 | 0.85 |

**Figure 4:**
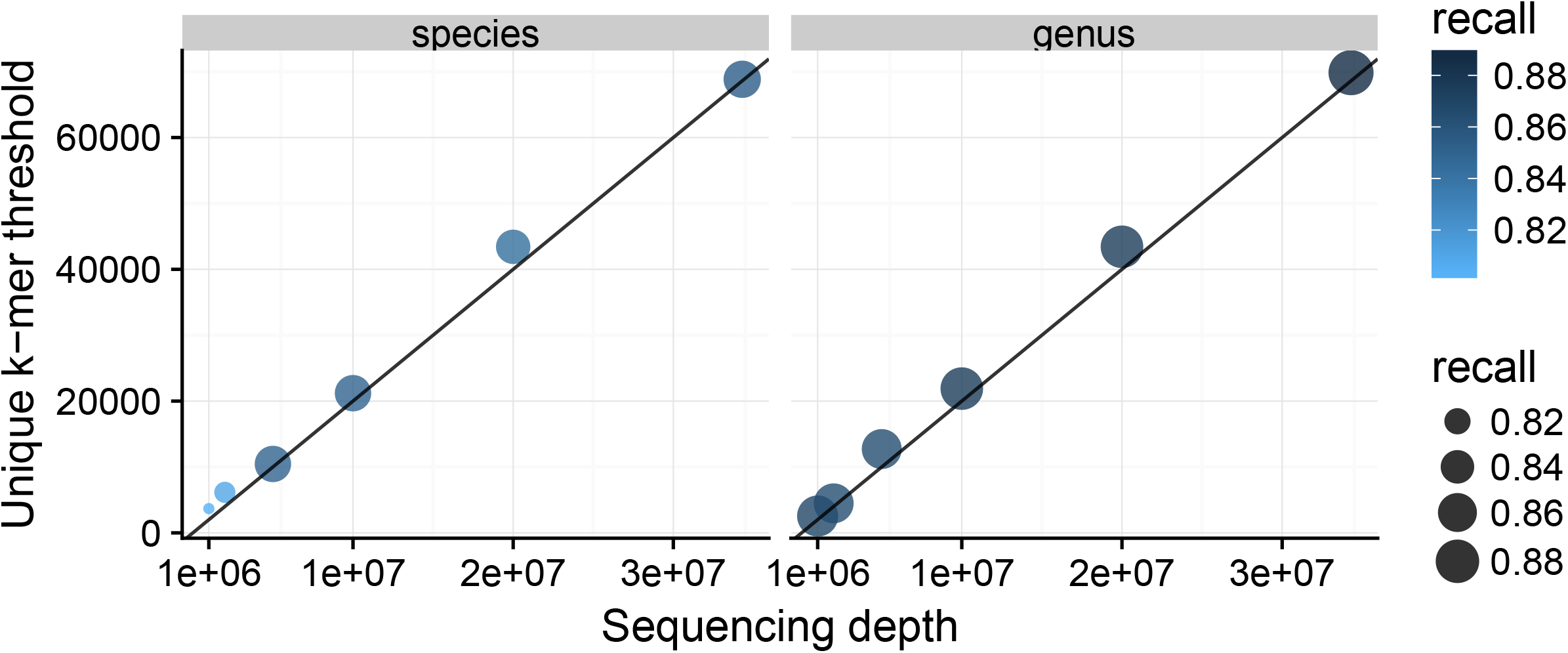
Deeper sequencing depths require higher unique k-mer count thresholds to control false-positive rate and achieve the best recall. A minimum threshold of about 2000 unique k-mer per a million reads gives the best results in this dataset (solid line in plot).

In general, we find that for correctly identified species, we obtain up to approximately *L-k* unique k-mers per each read, where *L* is the read length, because each read samples a different location in the genome. (Note that once the genome is completely covered, no more unique k-mers can be detected.) Thus the k-mer threshold should always be several times higher than the read count threshold. For the discovery of pathogens in human patients, discussed in the next section, a read count threshold of 10 and unique k-mer count threshold of 1000 eliminated many background identifications while preserving all true positives, which were discovered from as few as 15 reads.

### Results on biological samples for infectious disease diagnosis

Metagenomics is increasingly used to find species of low abundance. A special case is the emerging use of metagenomics for the diagnosis of infectious diseases [27, 28]. In this application, infected human tissues are sequenced directly to find the likely disease organism. Usually, the vast majority of the reads match (typically 95-99%) the host, and sometimes fewer than 100 reads out of many millions of reads are matched to the target species. Common skin bacteria from the patient or lab personnel and other contamination from sample collection or preparation can easily generate a similar number of reads, and thus mask the signal from the pathogen.

To assess if the unique k-mer count metric in KrakenHLL could be used to rank and identify pathogen from human samples, we reanalyzed ten patient samples from a previously described series of neurological infections [4]. That study sequenced spinal cord mass and brain biopsies from ten hospitalized patients for whom routine tests for pathogens were inconclusive. In four of the ten cases, a likely diagnosis could be made with the help of metagenomics. To confirm the metagenomics classifications, the authors in the original study re-aligned all pathogen reads to individual genomes.

Table 3 shows the results of our reanalysis of the confirmed pathogens in the four patients, including the number of reads and unique k-mers from the pathogen, as well as the number of bases covered by re-alignment to the genomes. Even though the read numbers are very low in two cases, the number of unique k-mers suggests that each read matches a different location in the genome. For example, in PT8, 15 reads contain 1570 unique k-mers, and re-alignment shows 2201 covered base pairs. In contrast, Table 4 shows examples of identifications from the same datasets that are not well-supported by k-mer counts. We also examined the likely source of the false positive identifications by blasting the reads against the full nt database, and found rRNA of environmental bacteria, human RNA and PhiX-174 mis-assignments (see Suppl. Methods for details). Notably, the common laboratory and skin contaminants PhiX-174, *Escherichia coli, Cutibacterium acnes* and *Delftia* were detected in most of the samples, too (see Suppl. Table 6). However, those identifications are solid in terms of their k-mer counts - the bacteria and PhiX-174 are present in the sample, and the reads cover their genomes rather randomly. To discount them, comparisons against a negative control or between multiple samples is required (e.g. with Pavian [29]).

**Table 3:** Validated pathogen identifications in patients with neurological infections have high numbers of unique k-mers per read. The pathogens were identified with as few as 15 reads, but the high number of unique k-mers indicates distinct locations of the reads along their genomes. Re-alignment of mapped reads to their reference genomes (column “Covered Bases”) corroborates the finding of the unique k-mers (see also Suppl. Figure 5). Interestingly, the k-mer count in PT5 indicates that there might be multiple strains present in the sample since the k-mers cover more than one genome. Read lengths were 150-250 bp.

| Sample | Matched microorganism | Reads | K-mers | Covered Bases |
| --- | --- | --- | --- | --- |
| PT5 | Human polyomavirus 2 | 9,650 | 7,129 | 5,130 / 5,130 |
| PT7 | *Elizabethkingia genomsp. 3* | 403 | 20,724 | 53,256 / 4,433,522 |
| PT8 | *Mycobacterium tuberculosis* | 15 | 1,570 | 2,227 / 4,411,532 |
| PT10 | Human gammaherpesvirus 4 | 20 | 2,084 | 2,822 / 172,764 |

**Table 4:** False positive identifications have few unique k-mers. Using an extended taxonomy, the identifications in PT4 and PT10 were matched to single accessions (instead of to the species level). The likely true source of the mapped sequences was determined by subsequent BLAST searches and included 16S rRNA present in many uncultured bacteria, human small nucleolar RNAs (snRNAs), and phiX174.

| Sample | Matched microorganism | Reads | K-mers | Source |
| --- | --- | --- | --- | --- |
| PT3 | *Clostridioides difficile* | 122 | 126 | 16S rRNA |
| PT4 | Hepatitis C virus<br>JF343788.1 Recombinant Hepatitis C virus | 101 | 3 | Human<br>snRNA |
| PT5 | *Akkermansia muciniphila* | 936 | 136 | 16S rRNA |
| PT10 | Human betaherpesvirus 5<br>JN379815.1 Human herpesvirus 5 strain<br>U04, partial genome | 63 | 5 | phiX174 |

### Further extensions in KrakenHLL

KrakenHLL adds three further notable features to the classification engine.

1. Enabling strain identification by extending the taxonomy: The finest level of granularity for Kraken classifications are nodes in the NCBI taxonomy. This means that many strains cannot be resolved, because up to hundreds of strains share the same taxonomy ID. KrakenHLL allows extending the taxonomy with virtual nodes for genomes, chromosomes and plasmids, and thus enabling identifications at the most specific levels (see Suppl. Methods Section 3)
2. Integrating 100,000 viral strain sequences: RefSeq includes only one reference genome for most viral species, which means that a lot of the variation of viral strain is not covered in a standard RefSeq database. KrakenHLL sources viral strain sequences from the NCBI Viral Genome Resource that are validated as ‘neighbors’ of RefSeq viruses, which leads to up to 20% more read classifications (see Suppl. Methods Section 4).
3. Hierarchical classification with multiple databases. Researcher’s may want to include additional sequence sets, such as draft genomes, in some searches. KrakenHLL allows to chain databases and match each k-mer hierarchically, stopping when it found a match. For example, to mitigate the problem of host contamination in draft genomes, a search may use the host genome as first database, then complete microbial genomes, then draft microbial genomes. More details are available in Suppl. Method Section 5.

### Timing and memory requirements

The additional features of KrakenHLL come without a runtime penalty and very limited additional memory requirements. In fact, due to code improvements, KrakenHLL often runs faster than Kraken, particularly when most of the reads come from one species. On the test dataset, the mean classification speed in million base-pairs per minute increased slightly from 410 to 421 Mbp/m (see Suppl. Table 3). When factoring in the time needed to summarize classification results by kraken-report, which is required for Kraken but part of the classification binary of KrakenHLL, KrakenHLL is on average 50% faster. The memory requirements increase on average by 0.5 GB from 39.5 GB to 40 GB.

On the pathogen Id patient data, where in most cases over 99% of the reads were either assigned to human or synthetic reads, KrakenHLL was significantly faster than Kraken (Suppl. Table 5). The classification speed increased from 467 to 733 Mbp/m. The average wall time was about 44% lower, and the average additional memory requirements were less than 1GB, going from 118.0 to 118.4 GB. All timing comparisons were made after preloading the database and running with 10 parallel threads.

## Discussion

In our comparison, KrakenHLL performed better in classifying metagenomics data than many existing methods, including the alignment-based methods Blast [16], Diamond [30], and MetaFlow [19]. Blast and Diamond results were post-processed by Megan [31], which assigns reads to the lowest-common ancestor (LCA), but ignores coverages when computing the resulting taxonomic profile. Thus, the taxonomic profile (with read counts as abundance measures) is sensitive to over-representing false positives that have coverage spikes in parts of the genome in the same way as non-alignment based methods. Coverage spikes may appear due to wrongly matched common sequences (e.g. 16S rRNA), short amplified sequences floating in the laboratory, and contamination in database sequences. MetaFlow, on the other hand, implements coverage-sensitive mapping, which should give better abundance calls, but it did not perform very well in our tests. Going from alignments to a good taxonomic profile is difficult because coverage information cannot be as easily computed for the LCA taxon and summarized for higher levels in the taxonomic tree. In comparison, reads and unique k-mer counts can be assigned to the LCA taxa, and summed to higher levels. Notably, KrakenHLL’s k-mer counting is affected by GC biases in the sequencing data the same way as other read classifiers and aligners [32], and may underreport GC-rich or GC-poor genomes.

## Conclusions

KrakenHLL is a novel method that combines fast k-mer based classification with an efficient algorithm for counting the number of unique k-mers found in each species in a metagenomics dataset. When the reads from a species yield many unique k-mers, one can be more confident that the taxon is truly present, while a low number of unique k-mers suggests a possible false positive identification. We demonstrated that using unique k-mer counts provides improved accuracy for species identification, and that k-mer counts can help greatly in identifying false positives. In our comparisons with multiple other metagenomics classifiers on multiple metagenomics datasets, we found that KrakenHLL consistently ranked at the top. The strategy of counting unique k-mer matches allows KrakenHLL to detect that reads are spread across a genome, without the need to align the reads. By using a probabilistic counting algorithm, KrakenHLL is able to match the exceptionally fast classification time of the original Kraken program with only a very small increase in memory. The result is that KrakenHLL gains many of the advantages of alignment at a far lower computational cost.

## Declarations

### Availability of data and material

KrakenHLL 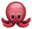 is implemented in C++ and Perl. Its source code is available at https://github.com/fbreitwieser/krakenhll, licensed under GPL3. The version used in the manuscript is permanently available under https://doi.org/10.5281/zenodo.1252385. Analysis scripts for the results of this manuscript are available at https://github.com/fbreitwieser/krakenhll-manuscript-code.

The datasets of McIntyre et al. are available at https://ftp-private.ncbi.nlm.nih.gov/nist-immsa/IMMSA. The sequencing datasets of Salzberg et al. are available under the BioProject accession PRJNA314149 (https://www.ncbi.nlm.nih.gov/bioproject/PRJNA314149). Note that human reads have been filtered. The test datasets generated by sampling reads from bacterial isolate SRA experiments are available at ftp://ftp.ccb.jhu.edu/pub/software/krakenhll/SraSampledDatasets.

## Acknowledgements

Thanks to Jen Lu, Ales Varabyou, Thomas Mehoke, David Karig, Sharon Bewick and Peter Thielen for valuable discussions on the general method and its applicability. Thanks to Alexa McIntyre and Rachid Ounit for providing very quick answers, scripts and data to their benchmarking paper. Thanks to Daniel Baker for suggestions on the HyperLogLog algorithm, and to Jessica Atwell for proofreading the manuscript.

## Authors’ contributions

FPB conceived and implemented the method. FPB and SLS wrote the manuscript. All authors read and approved the final manuscript.

## Ethics approval and consent to participate

Not applicable.

## Consent for publication

Not applicable.

